# A genome language model for mapping DNA replication origins

**DOI:** 10.64898/2026.01.29.702604

**Authors:** Pauline L. Pfuderer, Francisco Berkemeier, Joelle Nassar, Alastair Crisp, Jedrzej J. Jaworski, Jessica Moore, Julian E. Sale, Michael A. Boemo

## Abstract

Origin firing is a central process during DNA replication, but specific sequences defining replication origin usage have not been defined in human cells. Here, we show that a genome language model can accurately predict which sequences can act as an origin of replication, thereby enabling the fast and cost-effective creation of genome-wide replication origin maps. We fine-tuned a genome language model on the primary sequence of mapped human origins to establish ORILINX (ORIgin of replication Language-model Inference via Nucleotide conteXt) and found that it learns a rich representation of sequence features linked to replication initiation, extending beyond known predictive features such as GC-content and G-quadruplex motifs. When applied genome-wide, the model’s sequence-derived origin calling closely mirrors origin efficiency inferred from replication timing, suggesting that intrinsic sequence context encodes information relevant to initiation frequency. Furthermore, we performed Short Nascent Strand sequencing (SNS-seq) and Repli-seq to demonstrate that ORILINX can generalise to other mammalian genomes, such as those of mice and sheep, as well as other vertebrates such as chickens. Finally, we packaged ORILINX into a simple, easy-to-use tool which is available at https://github.com/Pfuderer/ORILINX.git.

## Introduction

To replicate the human genome in a timely manner, DNA replication is initiated at 30,000 to 50,000 sites throughout the genome, termed “origins of replication” [1, 2]. Unlike in the budding yeast *S. cerevisiae*, origins of replication in human cells are not thought to be sequence-specific, meaning that there is no pattern of nucleotides associated with sites that serve as origins of replication [3, 4]. However, studies have identified several genomic features that human origins of replication share: higher GC-content, and association with CpG islands, open chromatin conformation, and DNase I accessibility, and G-quadruplex (G4) structure forming motifs [2, 5–8]. However, these features are neither necessary nor sufficient for a locus to act as an origin [6, 9]. As a result, traditional motif-finders or k-mer-based approaches have failed to produce accurate *in silico* classifiers of mammalian replication origins.

Importantly, many of the organisational principles that govern DNA replication in human cells are conserved across mammals. Genome-wide studies in mouse and other species have revealed broadly conserved replication timing programmes, reflecting a reproducible temporal order in which large genomic domains replicate during S phase [10–12]. Replication timing, however, is a domain-scale property and does not imply one-to-one conservation of individual initiation sites. Instead, it is consistent with the idea that many mammalian genomes share common constraints on where origins are licensed and fired. In this view, human replication origins are a particularly well-characterised example of a broader mammalian strategy rather than a distinct case. This conservation motivates the hypothesis that determinants of origin competence, the ability of a locus to become licensed via MCM loading in G1, are at least partly encoded in primary sequence features that are shared across species, even if they do not manifest as simple, strictly conserved motifs [13, 14].

Genome language models (gLMs) offer new opportunities to address this challenge. These models are trained to understand the “language” of genomes in a manner analogous to large language models (LLMs) trained on natural language. gLMs are particularly well suited to interrogate non-coding regulatory DNA - which constitutes 98% of the human genome - and have recently demonstrated strong performance across a variety of regulatory prediction tasks by capturing long-range dependencies inaccessible to classical sequence models [15–22].

We developed ORILINX (ORIgin of replication Language-model Inference via Nucleotide conteXt), a gLM fine-tuned on experimentally mapped human origins of replication to identify origins of replication directly from primary sequence. ORILINX can identify genomic loci that act as efficient “core” origins, as well as weaker, context-dependent “stochastic” origins [23]. While we trained ORILINX on SNS-seq data from human cells, we performed further SNS-seq in chicken and sheep cell lines, as well as in a publicly available mouse cell line, to demonstrate that the zero-shot performance (or predictive accuracy on a target dataset the model was not trained on) of ORILINX in these organisms is comparable to its test performance in human cells. This suggests ORILINX can generalise across mammalian species, as well as more broadly to at least some other vertebrates. To further validate ORILINX, we performed Repli-seq on mouse, sheep, and chicken cells to use as input for our recently developed method that generates whole-genome mathematical models of replication from Repli-seq [24]. This showed a strong correlation between competent replication origins predicted by ORILINX and efficient sites of replication predicted by the model. To enable the community to predict replication origin landscapes at scale, we engineered ORILINX into a fast, accessible, and easy-to-use tool that only requires an indexed FASTA file as input (https://github.com/Pfuderer/ORILINX.git).

## Results

### ORILINX makes accurate origin predictions across the human genome

ORILINX is built upon the publicly available DNABERT-2 model [25] using the parameter-efficient Low Rank Adaptation (LoRA) method [26] and flash attention [27] for fine-tuning (Figure 1a). We used a set of 64,148 core origins, identified via Short Nascent Strand sequencing (SNS-seq) across six human cell lines for model training (Figure 1b) and evaluation [23]. The model was trained on 2-kilobase sequences centred on the SNS-seq peak midpoints (Supplementary Figure S1). The model therefore makes a prediction, in the form of a probability, at each 2-kilobase window of the input genome indicating whether that window is a competent replication origin. ORILINX was tuned such that we consider a probability above 0.5 to be a positive call.

**Fig. 1.**
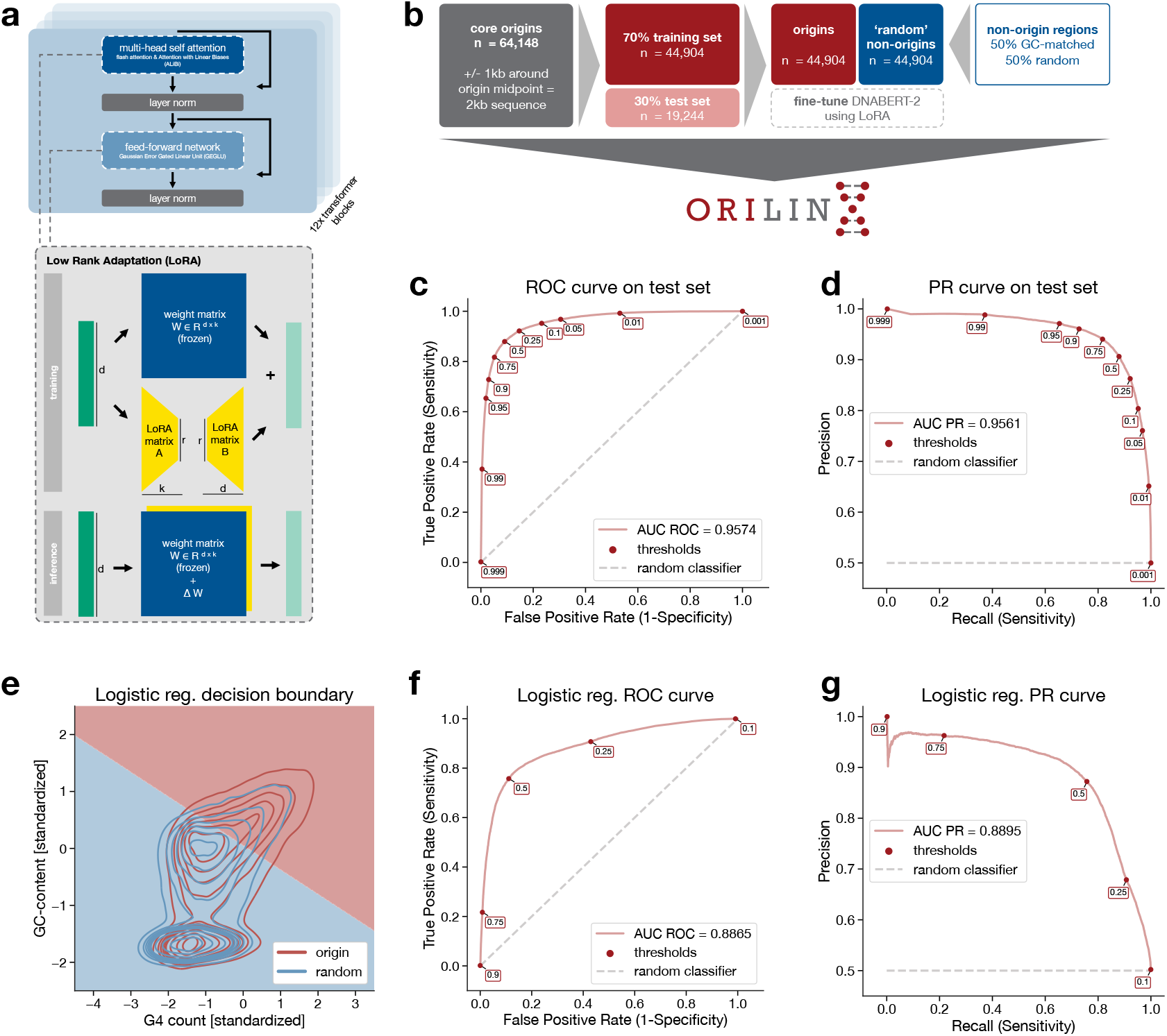
ORILINX model fine-tuning and evaluation on human core origins. **a)** Fine-tuning with parameter efficient Low Rank Adaptation (LoRA). **b)** Preparation of training data using the SNS-seq core origins set from [23]. **c)** Receiver operating characteristic (ROC) curve showing the ORILINX model performance on the 30% hold-out test set and matched number of random non-origin sequences, resulting in an area under the curve (AUC) of AUC ROC = 0.96. **d)** Same data as in **b)** showing the model performance in a Precision-Recall (PR) curve, reaching an AUC PR = 0.96. **e)** Kernel density estimate (KDE) plot showing the distribution in the test data of origin (red) and random non-origin (blue) sequences’ GC-content (y-axis, standardised) and G4 count (x-axis, standardised) alongside the learned linear regression boundary line. The logistic regression model was trained on the same training data and evaluated on the same test data as the fine-tuned DNABERT-2 model. **f)** ROC curve showing the logistic regression model performance on the 30% hold-out test set and matched number of random non-origin sequences (same data as in **c)** and **d)**), resulting in an AUC ROC of 0.89. **g)** Same data as in **f)** showing the model performance in a PR curve, reaching an AUC PR of 0.89.

We evaluated the model on a test set comprised of 30% of the core origins (n = 19,244 origins with a matched number of genomic background sequences) which was not part of the training process. This resulted in an area under the curve (AUC) of a receiver operating characteristic (ROC) curve of 0.96 and an AUC of a precision-recall (PR) curve of 0.96 (Figure 1c,d; Supplementary Figure S2). To show that ORILINX was basing its predictions on novel features beyond GC-content and G4-count, we created a two-dimensional representation of the final layer embeddings to show that ORILINX does not merely partition sequences by GC-content and G4-count (Supplementary Figure S3). Furthermore, we created a simple logistic regression classifier based on GC-content and G4-count (Figure 1e) and found that our model outperformed the logistic regression by a wide margin as measured by both AUC ROC and AUC PR (Figure 1f,g), particularly when applied to weaker, cell-type specific stochastic origins (Supplementary Figure S4). Finally, to establish that our training procedure was independent of the underlying assay for identifying origins of replication, we re-trained an ORILINX model purely on Ini-seq2 data [6], achieving an AUC ROC of 0.99 and AUC PR of 0.98, illustrating that ORILINX does not rely on potential artefacts from SNS-seq to make positive origin calls (Supplementary Figure S8). Taken together, these data show that ORILINX can make highly accurate predictions of genomic loci for both core and stochastic human replication origins by using features of the primary sequence beyond simple GC-content and G4 motifs.

### Validation of ORILINX on well-studied human replication origins

To further validate the predictive power of ORILINX, we benchmarked it on four well-studied human origins located in *c-MYC*, in the promoter of *TOP1*, near the 3’ end of *LMNB2*, and in *HBB*. The human *c-MYC* locus contains an initiation zone spanning its upstream enhancer that binds the c-Myc protein and fires in the early S phase [28–30]. In contrast, firing at the *β*-globin (*HBB*) locus is developmentally regulated; while most somatic cells initiate *HBB* replication late in S phase, erythroid lineage cells initiate early [31], likely driven by the open chromatin state [32, 33]. Finally, *LMNB2* and *TOP1* contain highly efficient origins associated with their downstream and upstream promoters, respectively [34–36].

To ensure there was no data leakage from the training process when predicting origin probabilities in the human genome, we created a separate training dataset for each of these four origins consisting of SNS-seq data that excluded the chromosome on which the respective origin is located. We verified that the model’s performance, when trained on each of these four datasets, was comparable to the benchmarks shown in Figure 1 (Supplementary Figures S5-S6) and found that the ORILINX predictions for the *c-MYC, TOP1* and *LMNB2* origins overlap with the genomic positions that have been validated by both SNS-seq and Ini-seq2 (Figure 2a-c). The observation of lower origin prediction scores in the developmentally regulated *HBB* origin (Figure 2d) is in line with the observation that ORILINX experiences a drop in performance in cell-type-specific stochastic origins (Supplementary Figure S4) compared to core origins.

**Fig. 2.**
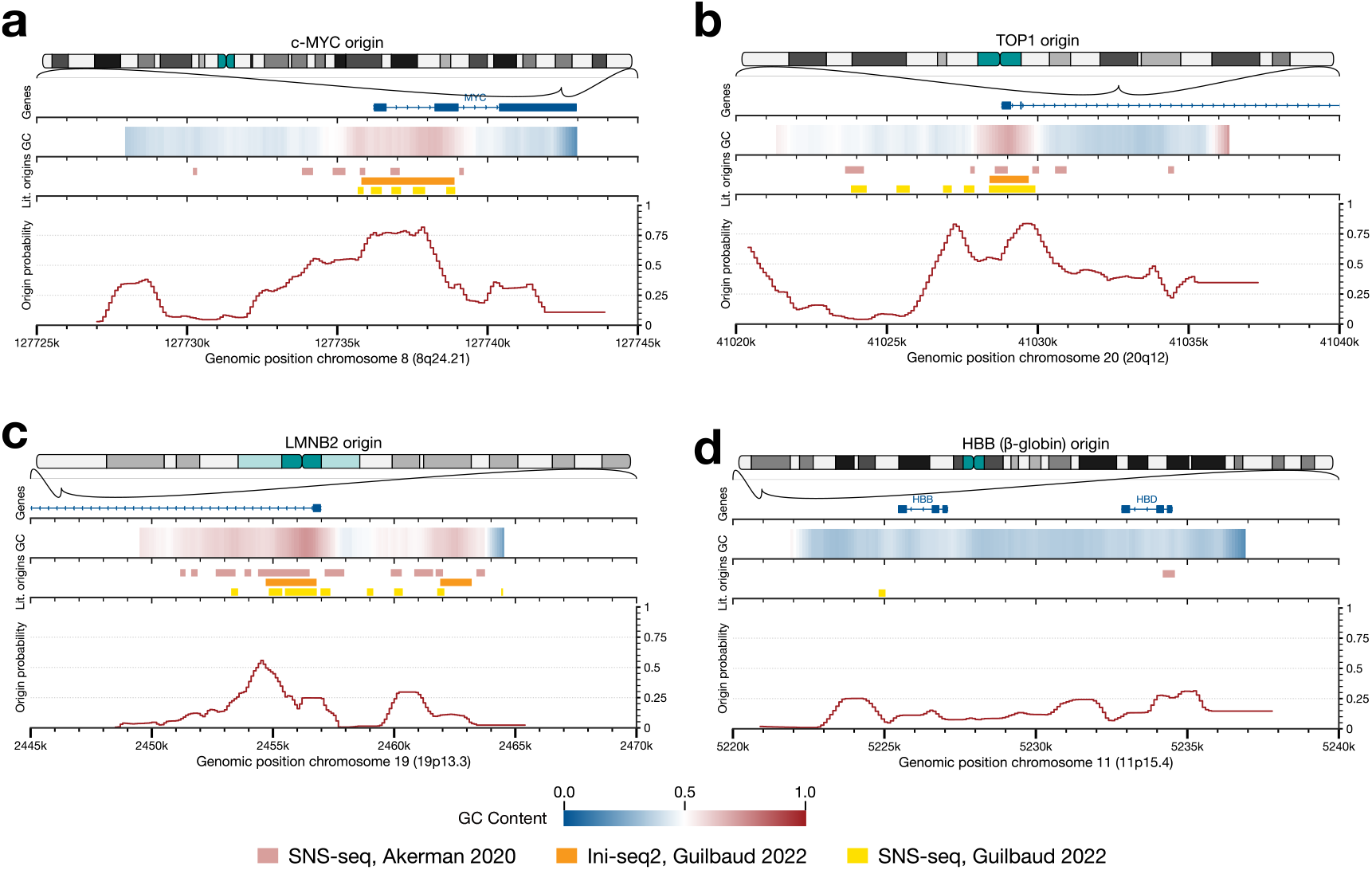
Origin predictions at well-studied human origins. Origin predictions with separately trained ORILINX chromosome-exclusion models for **a)** the *c-MYC* origin, predicted using a model where chromosome 8 was excluded from the training data **b)** the *TOP1* origin, predicted using a model where chromosome 20 was excluded from the training data, **c)** the *LMNB2* origin, where chromosome 19 was excluded from the training data and **d)** the *HBB* origin, where chromosome 11 was excluded from the training data. Each plot **(a-d)** shows the location of the locus on the respective chromosome, the GC content heatmap, known origins from the literature [6, 23], and the ORILINX predictions smoothed over 15 prediction windows of 2,000 bp each.

To juxtapose these four established replication origins, we used ORILINX to make predictions on loci that were established to be origin sparse. We investigated three long, late-replicating genes: *FHIT, NRXN1* and *WWOX* [24, 37]. Across all three of these regions, we observed low origin prediction scores (Supplementary Figure S7). Together, these results show that ORILINX learned sequence representations of human replication origins that enabled accurate prediction on chromosomes that it was not exposed to during training. The only input to ORILINX is primary sequence with no further information about cell type or developmental stage. As expected, ORILINX showed lower prediction scores at the developmentally regulated *β*-globin origin, consistent with the fact that the model relies exclusively on primary sequence features and does not incorporate cell-type-or stage-specific regulatory information. In contrast, the model performed well on the three established core origins in *c-MYC, TOP1* and *LMNB2*.

### Mathematical modelling shows that ORILINX predictions correlate with origin efficiency

ORILINX was trained and benchmarked on SNS-seq data and, while we also trained the model on Ini-seq2 data for the purpose of ensuring the model was not relying exclusively on SNS-seq artefacts, we sought an independent source of data to further validate ORILINX predictions. Replication timing, as measured by Repli-seq, indicates when genomic regions are replicated during S phase [38–40]. Previous mathematical modelling work has shown that, when assuming a constant rate of replication fork movement throughout the genome and that each origin can initiate stochastically with its own exponentially-distributed firing time, a genome-wide map of origin firing rates can be fit to a replication timing profile created from Repli-seq. Simulating these models makes it straightforward to estimate each origin’s efficiency, defined as the probability that an origin fires during a cell cycle [24, 41].

Following [24], we fit a whole-genome mathematical model to publicly available Repli-seq data from human H1 embryonic stem cells [40]. Forward simulation of the fitted model produced a genome-wide estimate of origin efficiency. We expected that regions with higher origin competence, defined here as the intrinsic, sequence-based potential to form a replication origin, would tend to correlate with higher origin efficiency, and therefore compared ORILINX predictions with efficiency estimates estimated from the mathematical model (Figure 3a). ORILINX makes predictions at each 2-kilobase window and, as shown in Figure 2, this high resolution means that the magnitude of the predictions can fluctuate even over relatively small regions. As we were interested in broad trends between the ORILINX predictions and the model, we used a 10-megabase moving-average window to smooth the ORILINX predictions which revealed a clear correlation between ORILINX prediction and the model’s estimate of efficiency (Spearman’s *ρ* = 0.89, p-value≪ 0.0001; Figures 3b-c). We further verified that this correlation was robust to the choice of moving-average window size (Supplementary Figure S9).

**Fig. 3.**
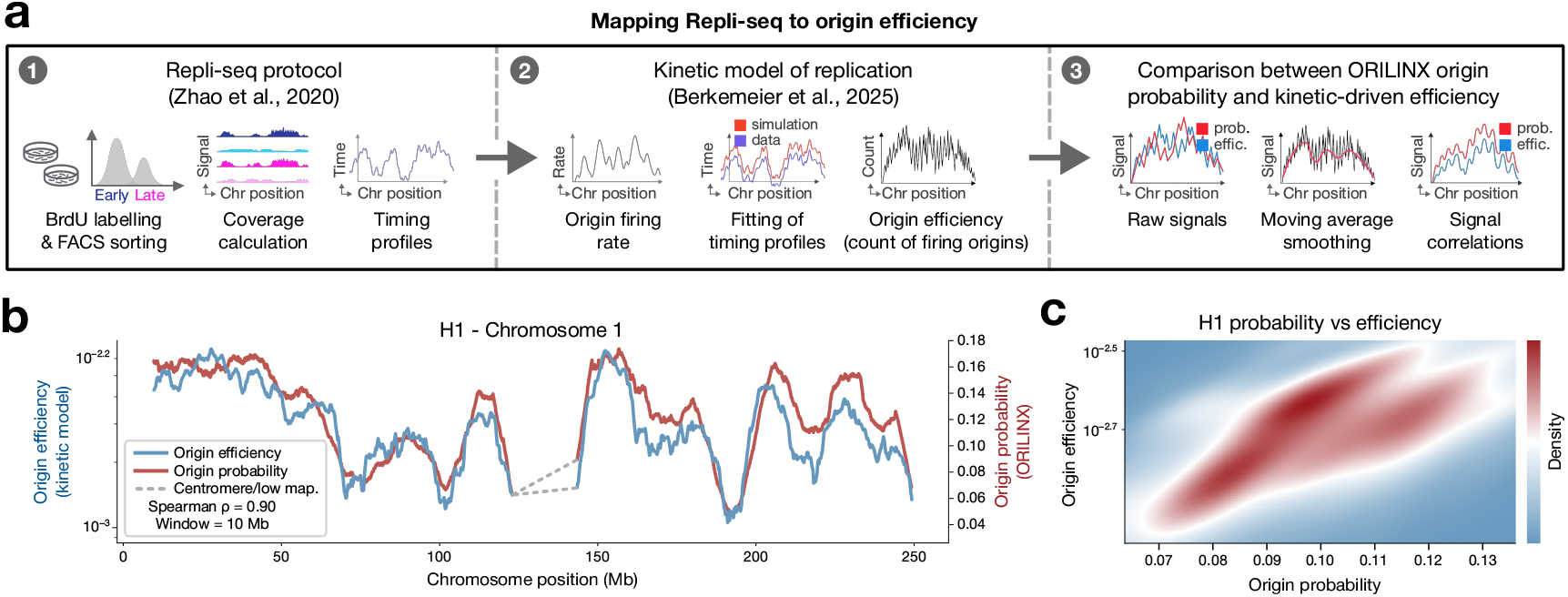
ORILINX origin probability correlates with replication-timing-inferred origin efficiency. **a)** Three-stage pipeline. (1) Experimental generation of a replication timing landscape via Repli-seq: BrdU incorporation during S phase, followed by Fluorescence-Activated Cell Sorting (FACS) into temporal fractions and sequencing-based quantification of replicated DNA, leading to a timing profile. (2) Computational inversion of the timing signal using a replication-kinetics model to estimate a genome-wide firing-rate landscape, from which origin efficiencies are computed directly from stochastic simulations governed by these rates. (3) Alignment of the inferred efficiencies with ORILINX sequence-derived origin probabilities for direct comparison on a common genomic scale. **b)** Comparison between ORILINX predicted origin probability (red) and timing-derived origin efficiency (blue) in chromosome 1 of the H1 human ES cell line, with both signals smoothed with a 10 Mb moving average window. Here, Spearman’s *ρ* = 0.90, p-value ≪ 0.0001. Centromere and low mappability regions are depicted with a grey dashed line and excluded from the correlation computation. **c)** Joint density distributions of probability and efficiency for whole-genome H1 cell line.

### Cross-species origin of replication site prediction with ORILINX and correlation with origin efficiencies

To demonstrate that ORILINX can generalise (Figure 4a) across mammalian genomes, we performed SNS-seq and Repli-seq on sheep primary fibroblasts and used previously published SNS-seq data from mouse embryonic stem cells [5]. Bench-marking ORILINX on the SNS-seq data showed that the performance on sheep fibroblasts was comparable to that of human cells shown in Figure 1c,d (AUC ROC of 0.93 and AUC PR of 0.94 on sheep fibroblasts compared to AUC ROC of 0.96 and AUC PR of 0.96 in human cells; Figure 4b). Moreover, this strong performance was consistent across biological replicates (Supplementary Figure S10). In mouse embryonic stem cells, ORILINX showed weaker benchmarks with AUC ROC of 0.81 and AUC PR of 0.85 (Figure 4c). However, these benchmarks still demonstrate that ORILINX can generalise to mammalian genomes beyond the human genome on which it was trained.

**Fig. 4.**
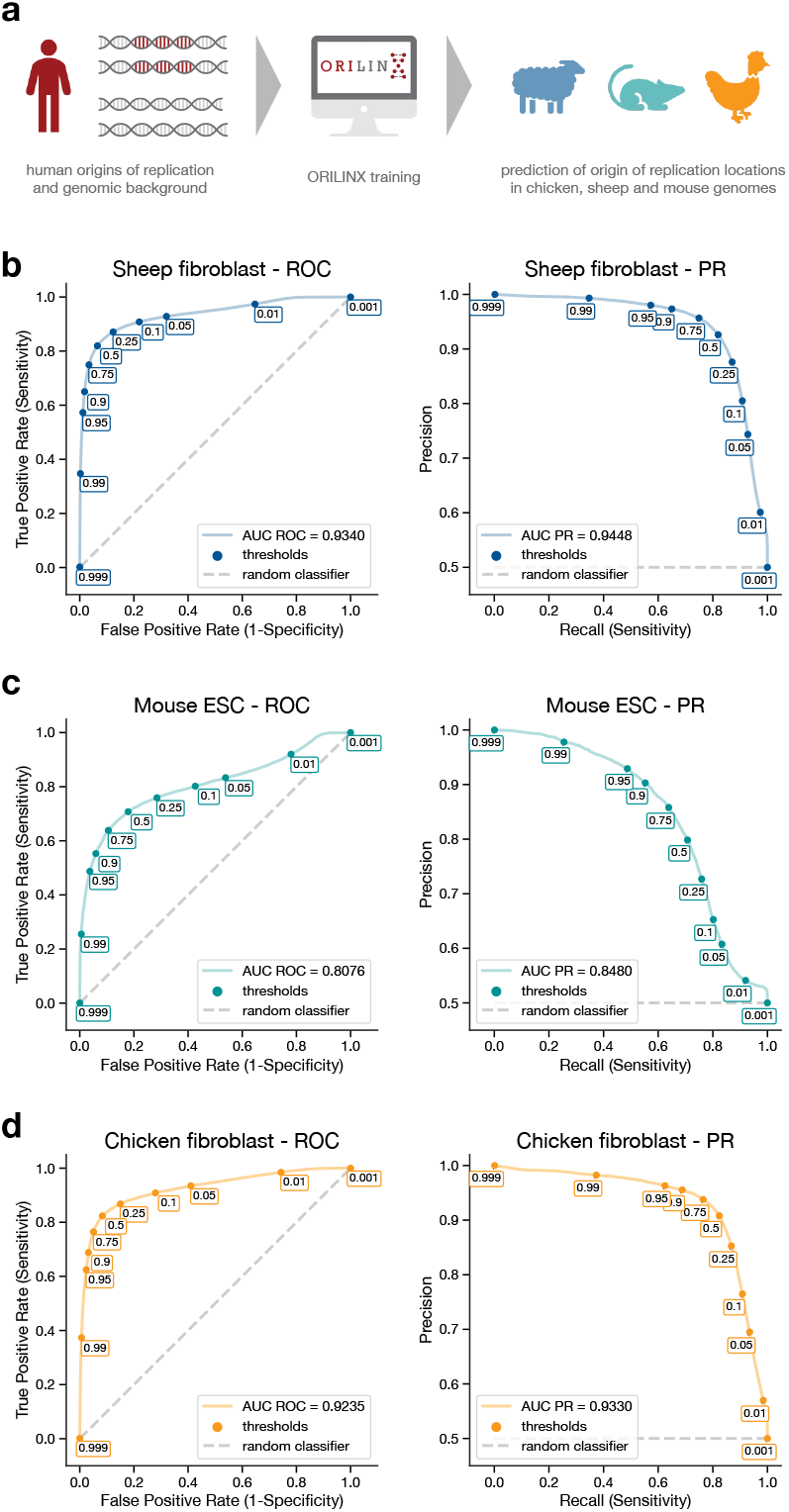
Cross-species ORILINX predictions. **a)** Schematic representation of ORILINX training on human origins of replication sequences, followed by applying ORILINX predictions in chicken, sheep and mouse without additional adjustments. **b)** ROC curves showing the ORILINX model performance in 28,490 SNS-seq chicken embryonic fibroblast cell origins of replication and matched number of random non-origin sequences, resulting in an AUC ROC = 0.92 and AUC PR =0.93. **c)** Same as in **b)** but for 79,574 SNS-seq sheep primary fibroblast origins and matched number random non-origin sequences from two replicates, resulting in an AUC ROC = 0.93 and AUC PR = 0.94. **d)** Same as in **b)** and **c)** but for publicly available mouse ESC data of 13,004 SNS-seq origins and matched number of random non-origin sequences, resulting in an AUC ROC = 0.81 and AUC PR = 0.85.

Next, we investigated whether ORILINX could generalise to other genomes beyond mammals. We therefore performed SNS-seq and Repli-seq on chicken embryonic fibroblast cells, where ORILINX achieved a zero-shot performance of AUC ROC 0.92 and AUC PR of 0.93 (Figure 4d). This performance is comparable to that in human cells and sheep fibroblasts, indicating that the sequence representations learned by ORILINX during training on the human genome are generalisable to at least some vertebrates beyond mammals.

Using our Repli-seq data for sheep primary fibroblasts, as well as Repli-seq data from [40] of mouse embryonic stem cells, we again constructed whole-genome mathematical models of genome replication for each organism and simulated these models to estimate each origin’s efficiency. As before, the correlation alignment between the ORILINX probability of origin competence and the mathematical model’s estimate of efficiency had a very strong correlation for both organisms (Figure 5).

**Fig. 5.**
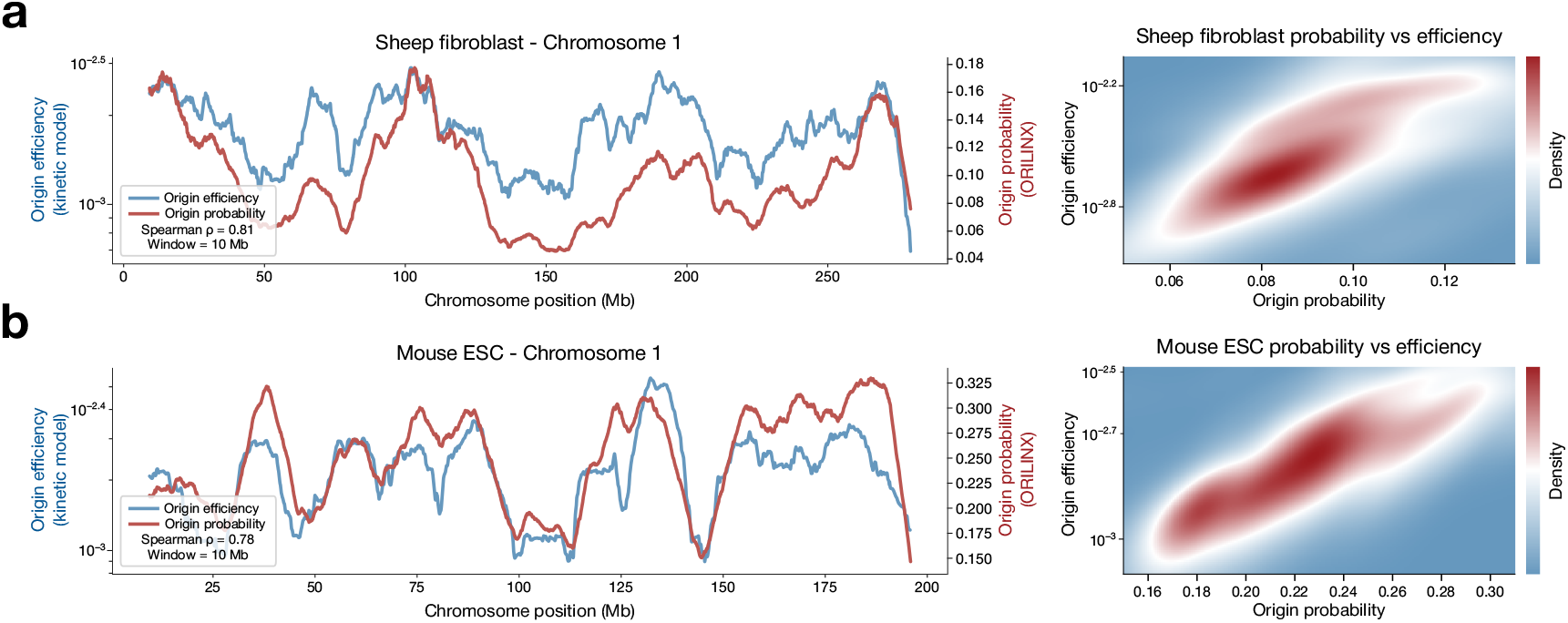
Cross-species origin efficiencies correlated with ORILINX predictions. Correlation analyses for **a)** sheep primary fibroblast, where Spearman’s *ρ* = 0.81, p-value ≪ 0.0001, and **b)** mouse embryonic stem cells (mESC), where Spearman’s *ρ* = 0.78, p-value ≪ 0.0001. For each species, the left panel shows chromosome 1 profiles comparing ORILINX predicted origin probability (red) with origin efficiency inferred from replication timing data (blue), with both signals smoothed using a 10 Mb moving average window. Spearman correlation coefficients are indicated. The right panel shows the corresponding joint density distributions of origin probability and origin efficiency computed genome-wide.

## Discussion

Here, we developed ORILINX, a fine-tuned gLM based on DNABERT-2 that can predict whether each 2-kilobase window along a genome is a competent replication origin. We showed that ORILINX predicts replication origins at well-studied loci, as well as a dearth of replication origins in long, late-replicating genes that are thought to be origin sparse. While ORILINX was trained exclusively on SNS-seq data from human cells, we demonstrated strong zero-shot performance on sheep and mouse cells, indicating that ORILINX can generalise well across mammalian genomes. Excitingly, ORILINX also showed strong zero-shot performance in chicken cells, suggesting that the model can generalise well to at least some other vertebrates. While ORILINX was trained, and largely benchmarked, on SNS-seq data, we used Ini-seq2 data to show that the model is not merely detecting SNS-seq artefacts, as well as Repli-seq and mathematical modelling to show that ORILINX tends to predict competent replication origins in genomic regions with a high number of efficient origins. Lastly, we engineered ORILINX into an efficient and easy-to-use tool that is available under the Apache 2.0 open-source license (https://github.com/Pfuderer/ORILINX.git).

ORILINX was designed to predict origin competence, but it is important to emphasise that these competence predictions are not independent of origin efficiency due to the way in which ORILINX was trained. Training on core origins, as defined by [23] (see Methods), means that ORILINX was trained to classify particularly efficient origins as competent. Loci were given binary labels rather than a measure of the origin’s activity from SNS-seq, hence we do not necessarily expect the ORILINX predictions to provide a measure of particular origin’s efficiency. Nevertheless, it stands to reason that regions with a particularly high number of competent origins would associate with both replication timing and origin efficiency, as shown in Figure 3 and Figure 5. The particular concordance between our mathematical models and ORILINX predictions raises the exciting opportunity of hybrid systems whereby ORILINX can be used to quickly inform and help generate whole-genome mathematical models of DNA replication in organisms for which no SNS-seq or Repli-seq data exists.

The one and only input to ORILINX is a primary sequence in FASTA format, typically a reference genome. This input carries no information about a particular cell type or development state, and hence this information is not taken into account at inference time. We therefore expect that ORILINX will perform well on core replication origins while showing a dip in performance on developmentally regulated replication origins, or indeed any origin whose firing is dependent on a particular cell type or state. This was reflected by lower averaged prediction scores around the *β*-globin origin (Figure 2d) as well as the performance on mouse embryonic stem cells which, while still strong, was not at the level of chicken or sheep fibroblasts (Figure 4d). For the mouse genome, however, an alternative (but not mutually exclusive) explanation comes from a particularly even distribution of origins compared to the human and chicken genomes [42]. This may make the mouse genome more challenging for ORILINX given the wider range of contexts. Nevertheless, it is encouraging that ORILINX outperformed simpler classifiers by a large margin, and that the relative improvement was wider on stochastic replication origins (Supplementary Figure S4).

ORILINX does not take any further information about cell type or state into account at inference time, but this information is still implicit in the data on which it is trained. ORILINX aims to predict the 2-kilobase windows which are capable of acting as replication origins and, should any developmentally regulated origins be missed, this could be alleviated with additional training data from the appropriate cell type, state, or stage. To that end, we envisage ORILINX growing considerably more accurate and comprehensive as more data continues to be generated.

We expect ORILINX to improve with further data, but at the same time, it is unrealistic to perform SNS-seq or other sequencing-based origin mapping techniques to every mammalian cell type. Here, we have shown that ORILINX can generalise remarkably well across mammals, despite being trained on a relatively small dataset. We therefore envisage ORILINX opening a fundamentally new door where, while its predictions may not be as accurate as performing SNS-seq, they are accurate enough that we can leverage the enormous difference in cost and scalability to generate a genome-wide origin map for every mammal with sequencing data available in a matter of hours. Moreover, and as we have shown, there is potential for ORILINX to generalise and scale to encompass more widely to birds or, more widely still, vertebrates.

## Methods

### Cell culture

Chicken embryonic fibroblast cells (UMNSAH/DF-1) were purchased from ATCC (ATCC-CRL-3586) and were maintained according to the supplier’s instructions in DMEM (ATCC 30-2002) supplemented with 10% fetal bovine serum (Gibco, 10270-106). Sheep primary fibroblasts were kindly provided by Madeline Lancaster (MRC Laboratory of Molecular Biology) [43], and maintained in DMEM GlutaMAX (Gibco, 31966-021) supplemented with 20% fetal bovine serum (Gibco, 10270-106), 50 µM 2-mercaptoethanol (Gibco, 21985-023) and antibiotics. All cells were grown at 37 °C in 5% CO_2_ and maintained between 30% and 90% confluency.

### Repli-seq

Repli-seq was performed as previously described [39]. Briefly, 5 to 10 ×10^6^ cells were pulsed with 100 µM BrdU (Sigma-Aldrich, cat.no B5002) for 2 h. Cells were then washed, collected, fixed by dropwise addition of ice-cold 70% ethanol, and stored at -20 °C until further processing. Samples were then treated with RNAse A (Thermo scientific, R1253) for 30 min and stained with Propidium Iodide (Invitrogen, P3566) before being sorted into four fractions corresponding to G1, early S (S1), late S (S2) and G2 phases, with 120,000 cells collected per fraction. Sorted cells were lysed in SDS-Protein K buffer for 2 h, followed by genomic DNA extraction (Zymo Quick-DNA microprep, D3020). Extracted DNA was sonicated to an average fragment size of 200 bp (Covaris M220: 175W, 10% duty cycle, 200 cycles per burst, 120 s, 4 - 7 °C water bath temperature). Sonicated DNA was subjected to end-repair and adaptor ligation using NEB-Next Ultra II DNA Library Prep Kit for Illumina (E7103). Adapter ligated DNA was then denatured and incubated with anti-BrdU antibody (12.5 µg/mL; BD cat. no. 555627), followed by immuno-precipitation with Rabbit anti-mouse IgG (20µg; Sigma-Aldrich, cat. no. M7023). Immunoprecipitated DNA was resuspended in digestion buffer with freshly added proteinase K. DNA was purified using DNA Clean & Concentrator-5 kit (Zymo Research, D4013). Final libraries were prepared using the NEBNext Ultra DNA Library Prep Kit for Illumina and sequenced on an Illumina NextSeq2000 (50bp, single end).

### SNS-seq

Short nascent strands were isolated as previously described [23], with minor modifications. Briefly, 2 ×10^8^ asynchronous cells were harvested and lysed using DNAzol reagent (Invitrogen, Cat. no.10503027). Genomic DNA was extracted and denatured prior to size fractionation by centrifugation through a 5-30% sucrose gradient (24,000 ×g for 20 h at 4 °C, no brake). Gradients were collected and analysed by alkaline agarose gel electrophoresis, and fractions containing DNA fragments of 1 to 2kb were pooled. Samples were subjected to two overnight λ exo digestions (150 U each, Custom order, Thermo Scientific), followed by DNA cleanup. Resulting ssDNA was subsequently converted into dsDNA by random priming (Invitrogen, cat. no. 18094-011). For control samples, the protocol was modified as follows. Instead of using treated genomic DNA as control, the 1-2 kb nascent strand sample was split into two equal aliquots. One aliquot constituted the experimental sample and was treated with λ exonuclease alone, whereas the second aliquot constituted the control sample and was treated with an RNAse A/XRN1 mix (Thermo scientific, R1253/New England Bio-labs, M0338) followed by *λ* exonuclease. This strategy was used to control for potential *λ* exonuclease processivity bias. Final libraries were prepared using NEBNext Ultra DNA Library Prep Kit for Illumina (E7103) and sequenced on an Illumina NextSeq2000 (50bp, single end).

### Origin dataset analysis

For the new SNS-seq analyses performed for this study, sequencing reads were quality trimmed using trim_galore (v0.6.10) with default parameters, then aligned to the relevant genome (sheep: Ramb_v2.0; chicken: galGal6) using bowtie2 (v2.5.1) with the following parameters: –end-to-end –sensitive -N 0. Peaks were called using macs2 (v2.2.9.1) with default parameters. The number of reads mapping to each peak for both sample and control was determined and the signal normalised by subtracting the control coverage from the sample coverage. Peaks were then ranked by normalised coverage and the top 20% of peaks taken as the Q1 and Q2 origins (as described in [23]). For Repli-seq analysis initially sequencing reads were quality trimmed using trim_galore (v0.6.10), then aligned to the relevant genome using bowtie2 (v2.5.1) with the following parameters: –end-to-end –sensitive -N 0. Duplicate reads were then removed using gatk MarkDuplicates (v4.3.0.0) with default parameters. Bigwigs were calculated using deeptools bamCoverage (v3.5.5) with the following parameters : – binSize 20000 –normalizeUsing RPKM. The datasets of human core and stochastic origins were taken from a previous study [23] where SNS-seq was used on 19 human samples, covering 6 cell types of untransformed cell lines – including cord blood hematopoietic cells (HC), human embryonic stem cells (hESC) and human mammary epithelial cells (HMEC) – and three immortalised cell lines – all derived from HMEC cells [23]. Origins were assigned to 10 quantiles based on average activity and defined the set of the top 2 quantiles (Q1 and Q2) were defined as “core” origins (n = 64,148), representing origins which are active independent of the cell type [23]. The remaining seven quantiles (Q3 to Q10) are referred to as “stochastic” origins (n = 256,600), representing origins not conserved across all cell types [23]. The Ini-seq 2 origins were obtained from [6] and are available on GEO under GSE186675. The previously published mouse SNS-seq data was obtained from [5] is available on GEO under GSE68347. The SNS-seq peaks were filtered to only include Q1 and Q2 peaks, as described by [23]. We used a window of ± 1,000 bp around the midpoint of the origin coordinates derived by SNS-seq, resulting in a total sequence length of 2,000 bp. The window size ±1,000 bp was chosen to capture any potentially influential sequence patterns upstream and downstream of the origin midpoint, while ensuring that the sequence length could still be managed in the DNABERT-2 model fine-tuning step with the computational resources available through the University of Cambridge High Performance Computer (HPC). The set of 64,148 core origins was randomly split into 70% training data (n = 44,904 origins) and 30% test data (n = 19,244 origins). Midpoint annotation and random splitting of the core origins dataset into training and test sets was performed using a custom Python script (version 3.11.5). The 30% test data was kept separate from any steps in the training process (methods section 4.3.3). We trained the model on the core origins only to ensure the model was only exposed to origin sequences occurring in a variety of cell types as opposed to the stochastic origin dataset, which comprises cell-type specific origins. The resulting start and end coordinates for each origin in the respective core training and test and stochastic datasets are stored in csv files which are then loaded by a custom data loader module as part of the adapted ORILINX model. The sequence for each 2-kilobase interval was obtained using pysam (version 0.23.3) and the hg38 human reference genome (initial release version; downloaded from https://hgdownload.soe.ucsc.edu/goldenpath/hg38/bigZips/). To assess the number of potentially G4-forming sequences in the 2-kilobase input sequences, G4 scores for each sequence were obtained using G4Hunter [44] with a window size of 25 and a threshold of 1.5. To handle outliers in total potentially G4-forming patterns per sequence, raw G4 counts per sequence were log transformed.

### Model architecture and parameter-efficient fine-tuning

We fine-tuned the publicly available DNABERT-2 model (DNABERT-2-117M; Hugging-Face ID zhihan1996/DNABERT-2-117M, commit d064dece8a8b41d9fb8729fbe3435278786931f1) for origin of replication classification. The model was wrapped in a custom PyTorch module (DnaBertOrigin-Model) and loaded using the original configuration with trust_remote_code=True to enable the DNABERT-2 implementation of flash attention and ALiBi positional biases. Where available in the configuration, flash-attention flags (use_flash_attn, flash_attn, use_flash_attn_mha) were set to True to allow efficient long-sequence attention. The model returns hidden states and attentions for downstream interpretability analyses. Tokenisation followed the DNABERT-2 pretraining setup, using its 4,096-token Byte Pair Encoding (BPE) vocabulary on 2-kilobase windows centred on origin midpoints (see Data preprocessing section). Each sequence was encoded as variable-length tokens, prepended with a [CLS] token and terminated with [SEP] and [PAD] tokens as needed. Token IDs were mapped to 768-dimensional embeddings via the pretrained embedding matrix. To adapt DNABERT-2 to origin classification while keeping most parameters frozen, we employed Low-Rank Adaptation (LoRA) using the PEFT library (v0.10.0) [26]. LoRA adapters were inserted into both attention and feed-forward projections of the transformer blocks, targeting the modules Wqkv, dense, gated_layers and wo. We treated the task as sequence classification (TaskType.SEQ_CLS) and configured LoRA with rank r = 16, scaling α = 32, dropout 0.1 and bias=“none”. This configuration, selected from a grid search over learning rates (2 × 10^−5^ −5 × 10^−4^), LoRA ranks (4,8, 16, 32) and α values (8, 16, 32, 64), provided the best validation F1 while training 2.5% of the total parameters. A linear classification head (hidden size 768 → 1) was appended on top of the [CLS] embedding. Gradient check-pointing was enabled (gradient_checkpointing_enable()) to reduce memory usage during training. During the forward pass, the PEFT-wrapped DNABERT-2 model produced the last hidden state and hidden-state history; the [CLS] embedding was passed through the classifier to yield a single logit per sequence. Logits were used with PyTorch’s BCEWithLogitsLoss for binary origin versus non-origin classification.

### Training procedure and hyperparameter optimisation

Training data consisted of 2-kilobase windows centred on SNS-seq–defined origin midpoints and matched non-origin windows sampled from hg38 (see SNS-seq dataset and negative sampling sections). Datasets were loaded via a custom OriginClassificationDataset class, which also handled negative-set construction (50% GC-matched; 50% fully random, non-overlapping with origins, ≤10 ‘N’ bases, excluding chromosome Y). Labels were balanced (50% origin, 50% non-origin). All training and hyperparameter sweeps were run with batch_size = 8, sequence_length = 2000, and an 80/20 training–validation split (val_split_fraction = 0.2). We used AdamW as optimiser together with a linear learning-rate scheduler with warm-up, specified via scheduler_type=“linear” and a warm-up phase corresponding to 10% of total training steps (warmup_ratio = 0.1). Early stopping was implemented with a patience of 4 epochs, monitoring validation F1. We performed several SLURM array-based grid searches over the base learning rate (LR) from 5 ×10^−7^ up to 5 ×10^−4^, the LoRA *α* of 8, 16, 32 or 64 and corresponding LoRA rank r (= 0.5 * LoRA *α*) of 4, 8, 16 or 32. Each hyperparameter combination was trained for up to 20 epochs on the University of Cambridge HPC (single NVIDIA A100-SXM4 GPU with 80 GB VRAM, 32 CPU cores, 1.0 TiB RAM). For each run, the best epoch was selected according to maximum validation F1. The final ORILINX model used in all downstream analyses corresponded to LR = 2×10^−5^, LoRA rank r = 16, LoRA *α* = 32, with the best checkpoint obtained at epoch 6.

### Model evaluation

For final evaluation, we applied the selected ORILINX model (epoch 6 checkpoint) to a 30% hold-out test set of core origins (19,244 sequences) and an equal number of newly generated random non-origin sequences drawn without GC-matching. During inference, LoRA weights were merged with the frozen base weights to obtain the effective updated weight matrix, and logits were passed through a sigmoid function to obtain origin probabilities. Sequences with probability > 0.5 were classified as origins. The model achieved an accuracy of 0.89, precision of 0.91, recall of 0.88, F1 score of 0.89, ROC AUC 0.96 of and PR AUC of 0.96.

### Extraction of final layer’s embeddings

To assess the final layer’s embeddings to understand which features the model learned, the test set was analysed. The ORILINX model was loaded and the last hidden state tensors (one 768-dimensional vector for all T tokens) were saved as NumPy arrays (.npz). The [CLS] token embedding of each sequence was extracted and used as input for dimensionality reduction using Uniform Manifold Approximation and Projection (UMAP). Using UMAP, the 768-dimensions of each sequence were projected into a 2-dimensional space, using the umap-learn package (version 0.5.7) and setting n_neighbors = 15, min_dist = 0.1 and random_state = 42 for reproducibility. The embeddings extraction was limited to randomly sub-sampled 2,500 sequences per group, to keep the data size manageable and the visualisations clear.

### Logistic regression model

A logistic regression model was trained to predict origin of replication sequences based on two sequence features: GC-content and G4 count. The same set of training and test core origins was used as for the fine-tuning and evaluation of the ORILINX model. First, the features were scaled to unit variance using the scikit-learn (version 1.7.0) StandardScaler function. Then, scikit-learn’s LogisticRegression function was fitted on the core origins and negative samples training data set. Predictions were generated using the predict and predict_proba methods from the Pipeline function of scikit-learn. The logistic regression’s performance was assessed using AUC ROC and AUC PR.

### Mapping replication timing to origin firing rates and efficiencies

Our analysis of origin efficiency is based on the replication-timing model introduced in [24]. In this framework, each chromosome is discretised into 1-kilobase sites, and every site is treated as a potential origin of replication with an intrinsic firing rate *f*_*j*_. In other words, the time at which a given origin at site *j* fires is modelled as an independent exponential random variable with parameter *f*_*j*_. Once an origin fires, two replication forks emanate from that site and progress in opposite directions at a constant speed *v*, passively replicating downstream sites until they encounter forks from neighbouring origins or reach a chromosome end. Fork movement is assumed to be independent of origin firing. Under these assumptions, the replication time *T*_*j*_ of a genomic position *j* is defined as the minimum, over all potential origins *i*, of the firing time at *i* plus the deterministic travel time | *i*− *j* | */v* required for a fork to reach *j*. [24] derived a closed-form expression for the expected (or mean) replication time *E*[*T*_*j*_] given by

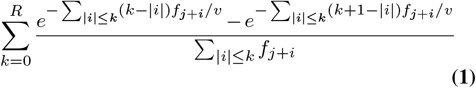

for a radius of influence *R*, i.e., the distance within which neighbouring origins are assumed to affect the timing of a focal origin. This expression is monotone in each *f*_*i*_, which makes it possible to invert the relationship between timing and firing. Given a genome-wide replication timing profile *T*_*j*_ obtained from Repli-seq, the model first constructs an initial estimate of the firing-rate landscape using an analytical approximation for the case of uniform firing, then iteratively updates each *f*_*j*_ to minimise the discrepancy between the predicted timing *E*[*T*_*j*_] and the experimentally measured *T*_*j*_. In practice, this iterative scheme converges rapidly and yields a unique firing-rate profile that reproduces the observed timing with low error.

Once the firing rates have been fitted, origin efficiencies follow naturally from the stochastic model. For each site *j*, the efficiency is defined as the probability that the origin at *j* fires before being passively replicated by a fork arriving from a neighbouring origin. This probability can be calculated either analytically from the fitted firing rates and constant fork speed, or numerically via stochastic simulations using a Gillespie-type algorithm. The resulting efficiencies provide a quantitative measure of functional origin usage across the genome and form a natural point of comparison for the sequence-based origin probabilities predicted by ORILINX, as discussed in Figure 3.

### Training exclusion controls confirm that correlations are not an artefact of genomic leakage

To ensure that the observed correlation between model-predicted origin probabilities and timing-derived origin efficiencies is not driven by inadvertent information leakage from training to evaluation, we performed a chromosome-exclusion control analysis. We trained three new ORILINX models, each one explicitly excluding all origin sequences from one chromosome during training: either chromosome 8, 11, 19 or 20. These chromosomes were selected arbitrarily and entirely withheld from the training dataset. When each model was evaluated on the held-out chromosome, the AUC ROC remained high, typically around 0.94 (Supplementary Figure S5), indicating that the classification performance does not rely on chromosome-specific memorisation, but instead reflects learned sequence principles that generalise across the genome.

For the same chromosome-excluded models, the genome-wide origin probabilities were then generated for the withheld chromosome, providing predictions made without any exposure to that chromosome’s origin sequences during model optimisation. A comparison between the non-leakage predictions and the original genome-wide probabilities showed that the model outputs are virtually identical (Supplementary Figure S6), indicating that leakage does not influence the predictions.

### Model and data availability

ORILINX is available at https://github.com/Pfuderer/ORILINX.git licensed under the Apache 2.0 open-source license. The pretrained DNABERT-2 model (DNABERT-2-117M) and pre-trained weights are available on HuggingFace (https://huggingface.co/zhihan1996/DNABERT-2-117M, commit version d064dece8a8b41d9fb8729fbe3435278786931f1). The Repli-seq and SNS-seq data for chicken and sheep are available at GEO accession code GSE317626. The publicly available SNS-seq core and stochastic origin coordinates are available under the original GEO accession code GSE128477 and the relevant two files are GSE128477_Core_origins_hg38.bed and GSE128477_Stochastic_origins_hg38.bed [23].

## ACKNOWLEDGEMENTS

This work was supported by a Cancer Research UK Cambridge Centre PhD Studentship awarded to P.L.P. (C9685/A25117) and J.M. (CTRQQR-2021/100012). This work was further supported by the Enrichment Programme at The Alan Turing Institute to P.L.P. F.B. and M.A.B. were supported by the Leverhulme Trust (RPG-2022-028) and the Biotechnology and Biological Sciences Research Council (UKRI1912). Additional funding and career support were provided by a Rokos Postdoctoral Associate position at Queens’ College Cambridge to F.B. and a fellowship at St John’s College, Cambridge to M.A.B. J.E.S., J.N., A.C. and J.J.J were supported by core funding to the MRC LMB from the Medical Research Council (MC_U105178808). J.J.J. was supported by Boehringer Ingelheim Fonds PhD Fellowship. This work was performed using resources provided by the Cambridge Service for Data Driven Discovery (CSD3), operated by the University of Cambridge Research Computing Service (https://www.csd3.cam.ac.uk), and supported by Dell EMC and Intel through Tier-2 funding from the Engineering and Physical Sciences Research Council (capital grant EP/T022159/1) and DiRAC funding from the Science and Technology Facilities Research Council (https://dirac.ac.uk).

## AUTHOR CONTRIBUTIONS

P.L.P. and J.M. conceived the project and P.L.P. led model development. F.B. developed the mathematical model. P.L.P. and M.A.B. developed the ORILINX tool. Model validation was performed by P.L.P., F.B. and J.J.J. Data were generated by J.N. and A.C. M.A.B. and J.E.S. supervised the study. The manuscript was written by P.L.P., F.B. and M.A.B. All authors read and approved the final manuscript.

## Supplementary Information

**Fig. S1.**
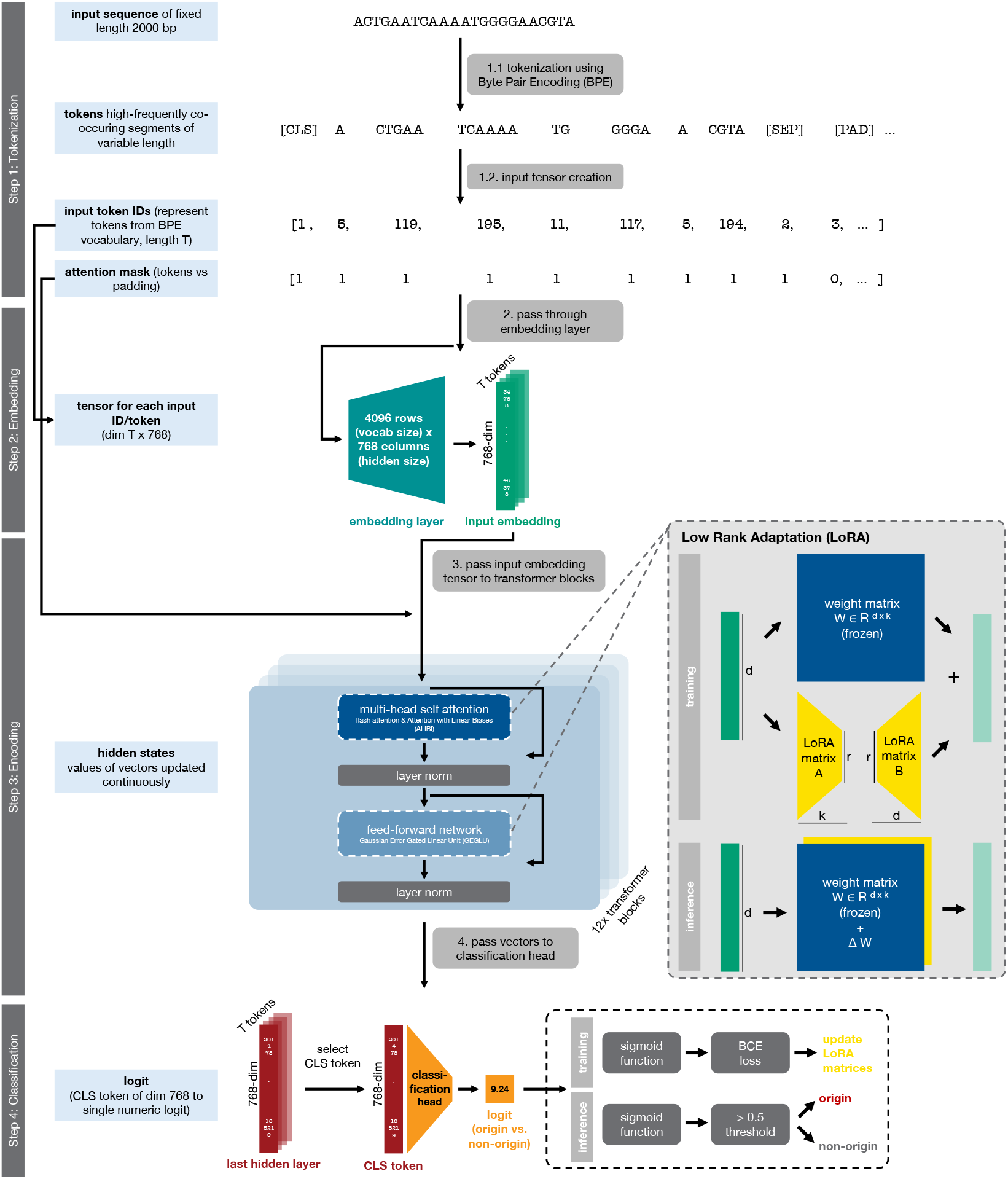
ORILINX model architecture and LoRA fine-tuning process. Step 1 (Tokenisation): The 2-kilobase DNA sequence centred around the origin midpoint is tokenized using Byte Pair Encoding (BPE) based on a previously established vocabulary of the most frequently co-occurring 4096 tokens. The input tensor is created by assigning each token a pre-defined numeric ID alongside an attention mask, indicating which positions are sequence or padding. The special classification token (CLS) contains information about all other sequence tokens, and the separation token (SEP) marks the end of the sequence, followed by padding tokens (PAD). Step 2 (Embedding): Each token is passed through the embedding layer, learned during the initial pre-training of DNABERT-2 [25], a 4096 × 768 dimensional matrix (teal coloured) resulting in each token being represented by a 768-dimensional input embedding vector (green). Step 3 (Encoding): The input tensor is passed through the BERT-style transformer blocks (12x blocks).The first step is a multi-head self-attention, including LoRA fine-tuning during the training step. Layer normalization is applied followed by a feed-forward network, again including LoRA fine-tuning in the training process. Step 4 (Classification): The CLS token is extracted from the last hidden layer and passed through a classification head, resulting in a logit, which is passed through a sigmoid function followed by binary cross entropy (BCE) loss calculation in the training process to inform LoRA matrix updates or thresholding (> 0.5) in the inference process, giving an indication of whether the input sequence is a origin sequence or not.

**Fig. S2.**
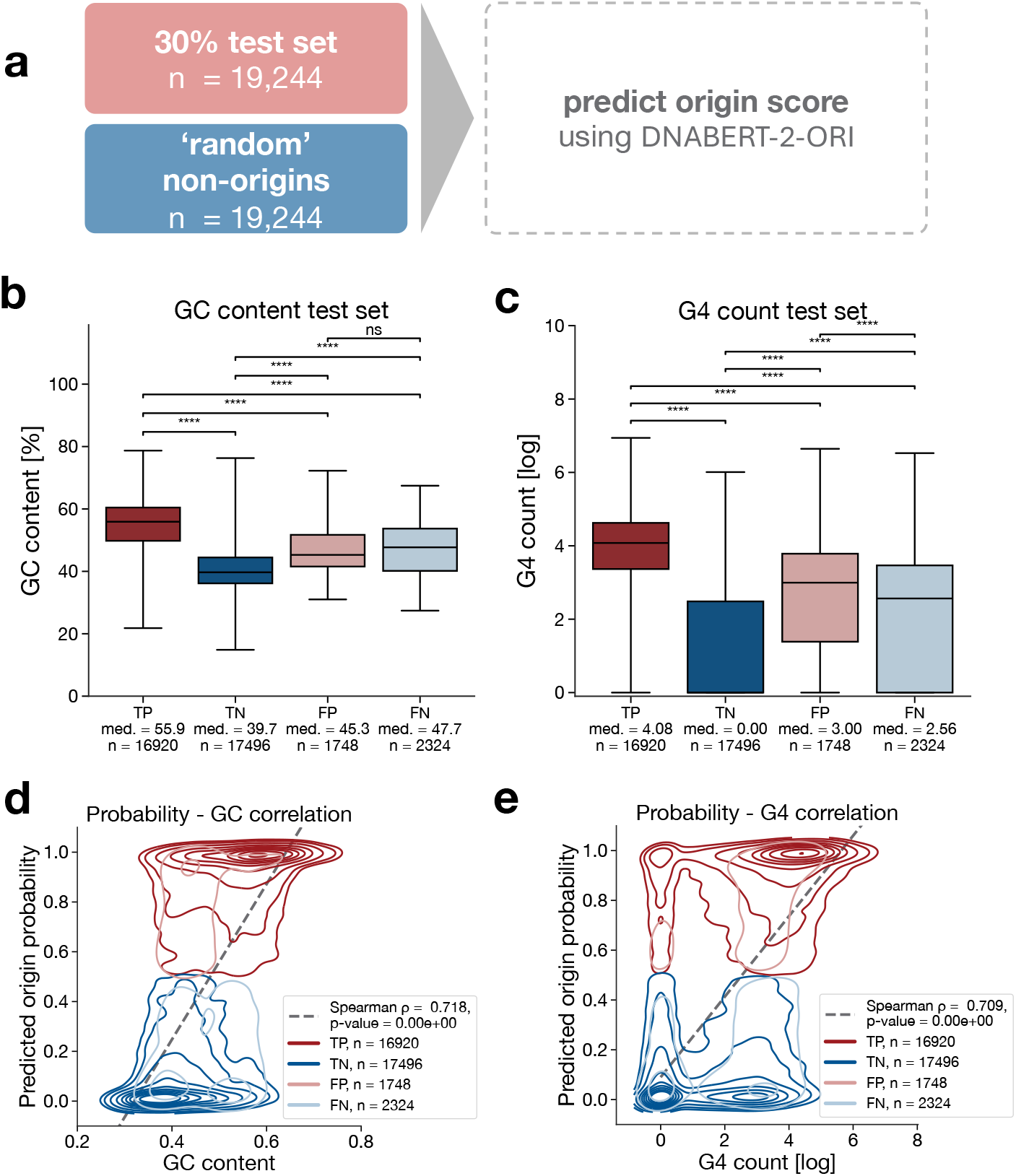
ORILINX model evaluation. **a)** The 30% hold-out test set of core origins from [23] used for model evaluation alongside newly generated random sequences (non GC-matched). **b)** Receiver operating characteristic (ROC) curve showing the ORILINX model performance on the 30% hold-out test set and matched number of random non-origin sequences, resulting in an area under the curve (AUC) of AUC ROC = 0.95. **c)** Same data as in **b)** showing the model performance in a Precision-Recall (PR) curve, reaching an AUC PR = 0.95. **d)** GC-content in the test set by classification result (true positive = TP, true negative = TN, false positive = FP, false negative = FN). **e)** G4 count in the test set split by TP, TN, FP and FN classification results. **f)** Correlation of GC-content and predicted origin probability by TP (dark red), TN (dark blue), FP (light red) or FN (light blue) group shown as kernel density estimates (KDEs). Overall correlation resulted in a Spearman’s *ρ* = 0.68, p-value ≪ 0.0001. **g)** Correlation of G4 count and predicted origin probability by TP (dark red), TN (dark blue), FP (light red) or FN (light blue) group shown as KDEs. Overall correlation resulted in a Spearman’s *ρ* = 0.69, p-value ≪ 0.0001. p-values for GC-content comparisons are obtained from a two-sided Welch’s t-test with no assumption of equal variances and p-values for G4 count comparisons are obtained from a two-sided non-parametric Wilcoxon Rank Sum test. Statistical significance: ns = not significant (p≥ 0.05), **** p ≪ 0.0001.

**Fig. S3.**
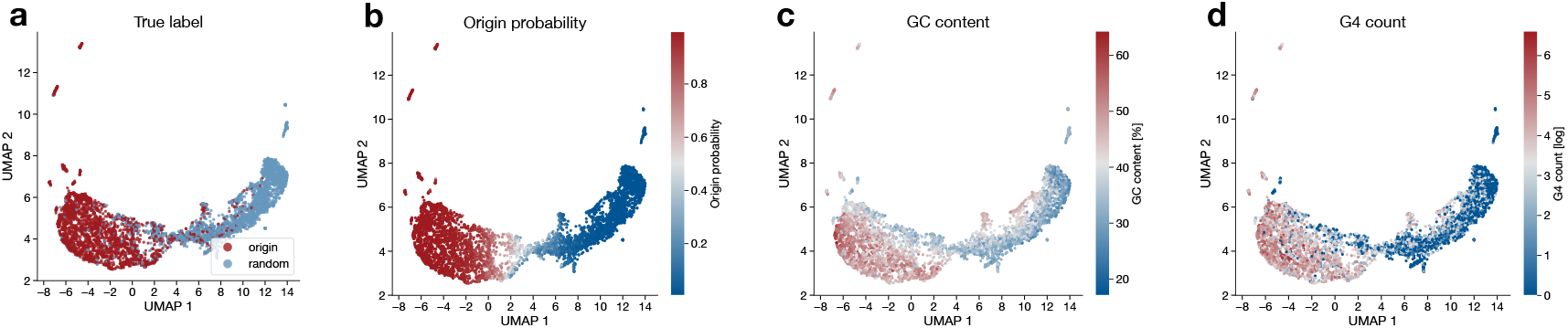
Final layer embeddings show separation beyond known origin of replication sequence features. 2-dimensional representation of the final 768-dimensional layer using UMAP. **a)** True sequence label. **b)** Inferred origin probability, with the threshold of calling a sequence an origin of replication of 0.5. **c)** GC-content percentage of across the 2,000 bp of each analysed sequence. **d)** Count of G4 sequences within each 2,000 bp long sequence. Note: 2500 randomly subsampled sequences each for origin or background sequences from the evaluation set of 19,244 sequences each.

**Fig. S4.**
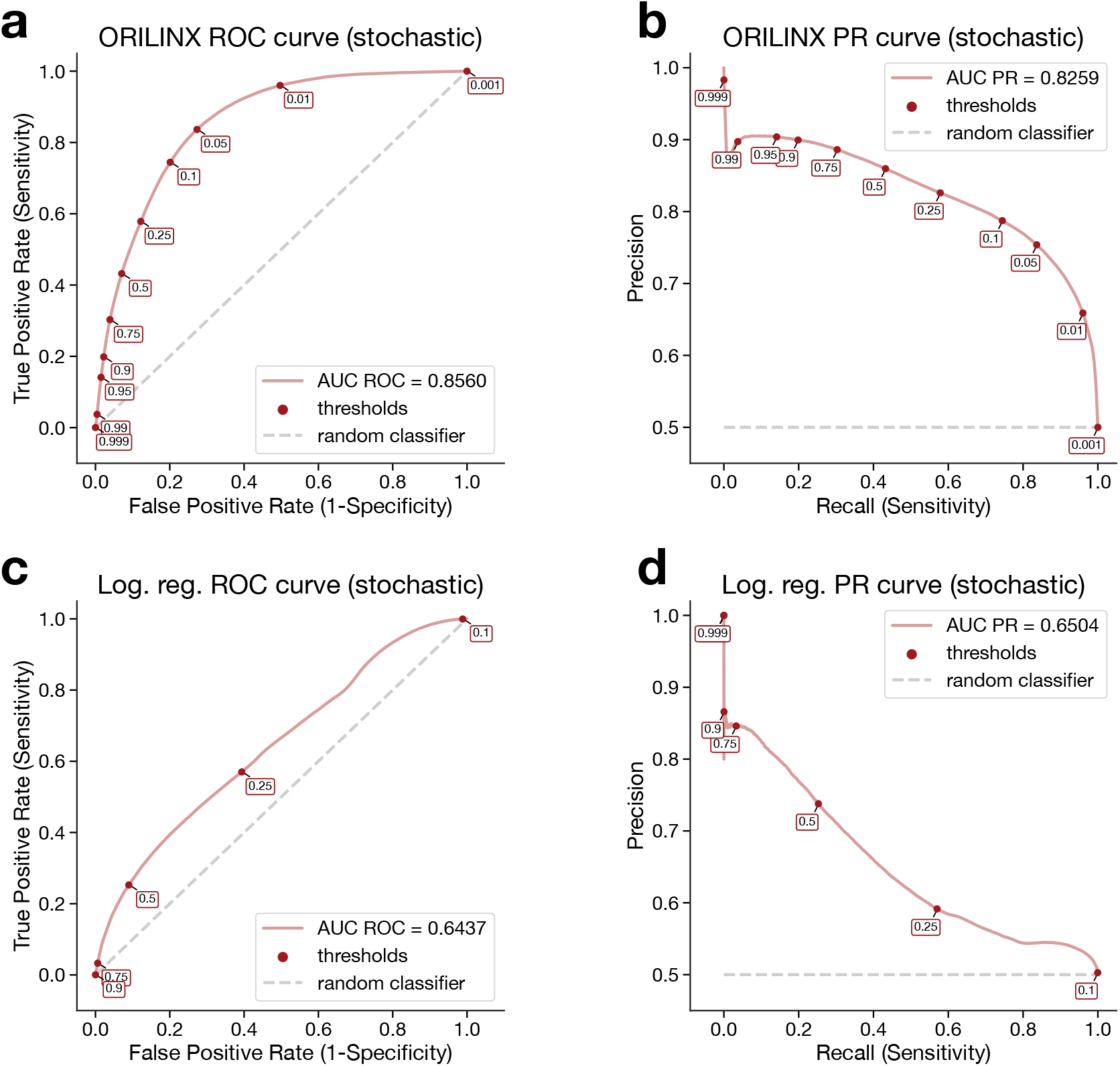
Generalisation of fine-tuned ORILINX or logistic regression model on the stochastic origins set. **a)** ROC curve for the fine-tuned ORILINX model when used for inference of the 256,600 stochastic origins [23] alongside a matched number of random non-origin sequences. **b)** PR curve for the fine-tuned ORILINX model tested on the stochastic origins data. **c)** AUC curve for the logistic regression model trained on the core origins and tested on the stochastic origins and **d)** PR curve for the logistic regression tested on the stochastic origins.

**Fig. S5.**
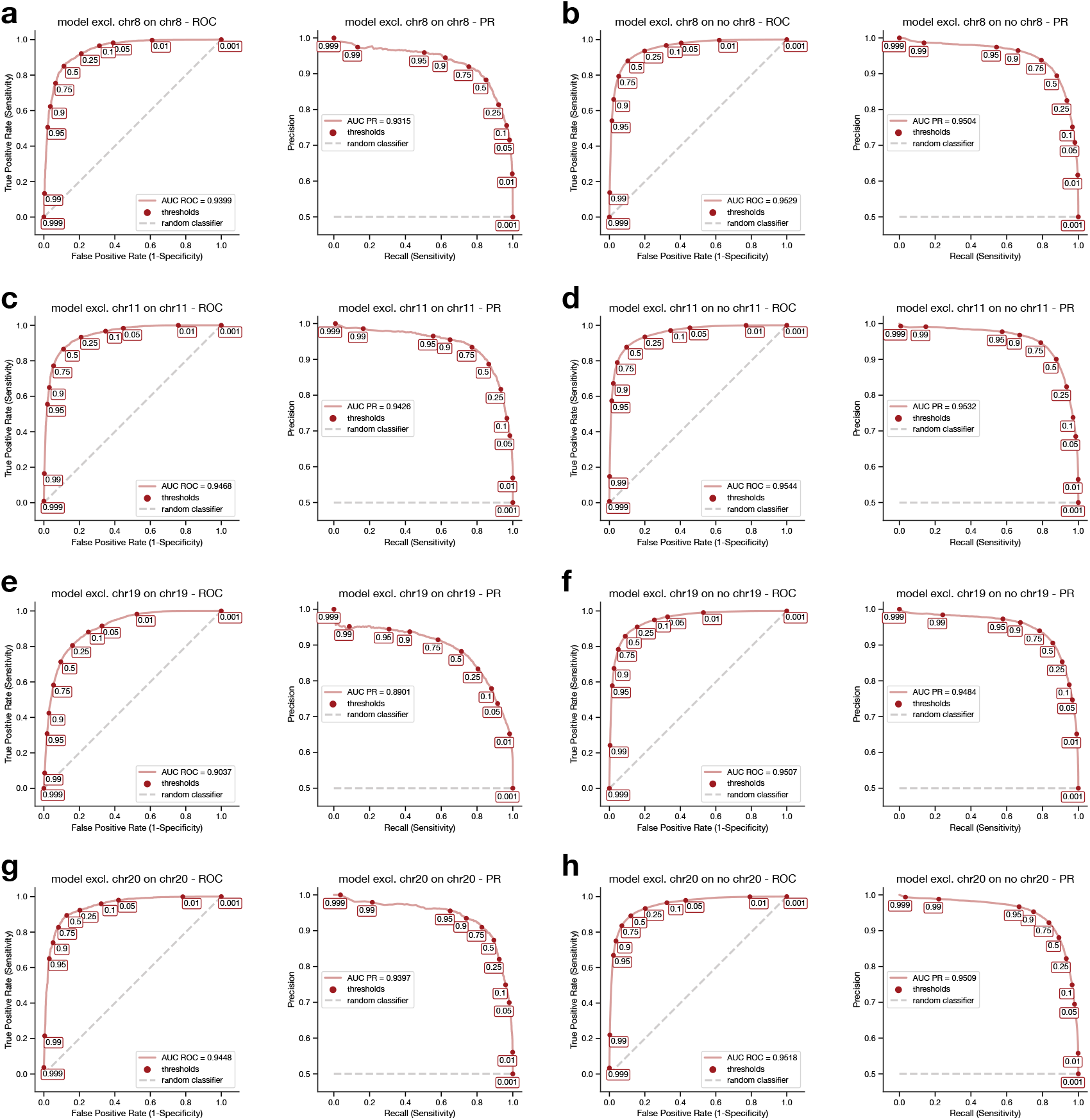
Performance for chromosome-exclusion ORILINX models. AUC ROC and AUC PR for models excluding the respective chromosome from training were evaluated on the respective chromosome test set **(a, c, e, g)** as well as on all remaining chromosomes test sets **(b, d, f, h)**.

**Fig. S6.**
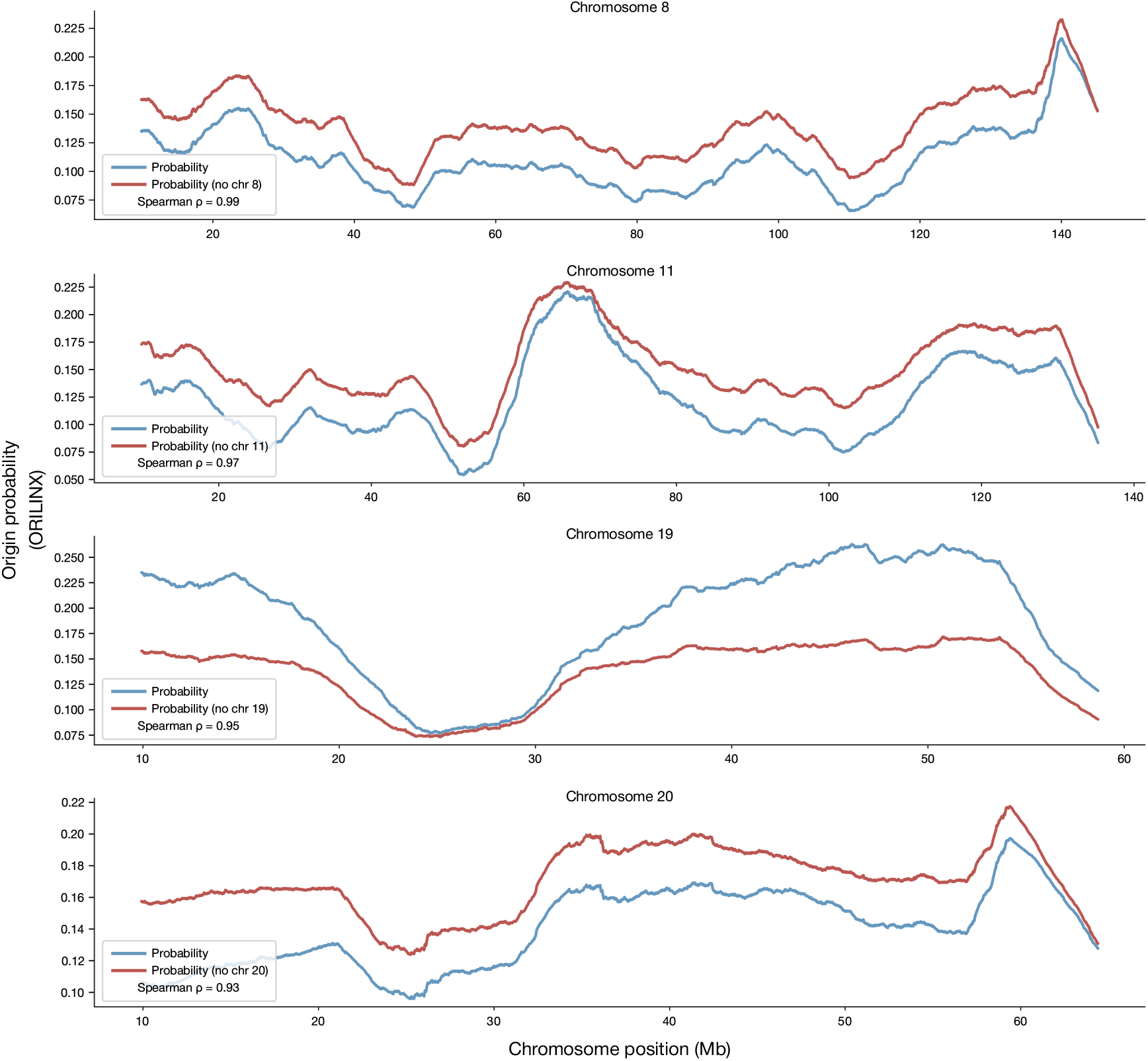
Comparison of ORILINX origin probabilities with and without chromosome-specific training leakage. Smoothed origin-probability tracks along human H1 chromosomes 8, 11, 19, and 20, comparing the standard ORILINX predictions (blue) with predictions obtained after excluding the corresponding chromosome from training (“no chr n”, red). Curves are plotted after a 10 Mb moving-average smoothing. Spearman correlation coefficients (*ρ*) between the two probability signals exceed 0.9 in all cases, confirming that ORILINX predictions are robust to potential training-set leakage.

**Fig. S7.**
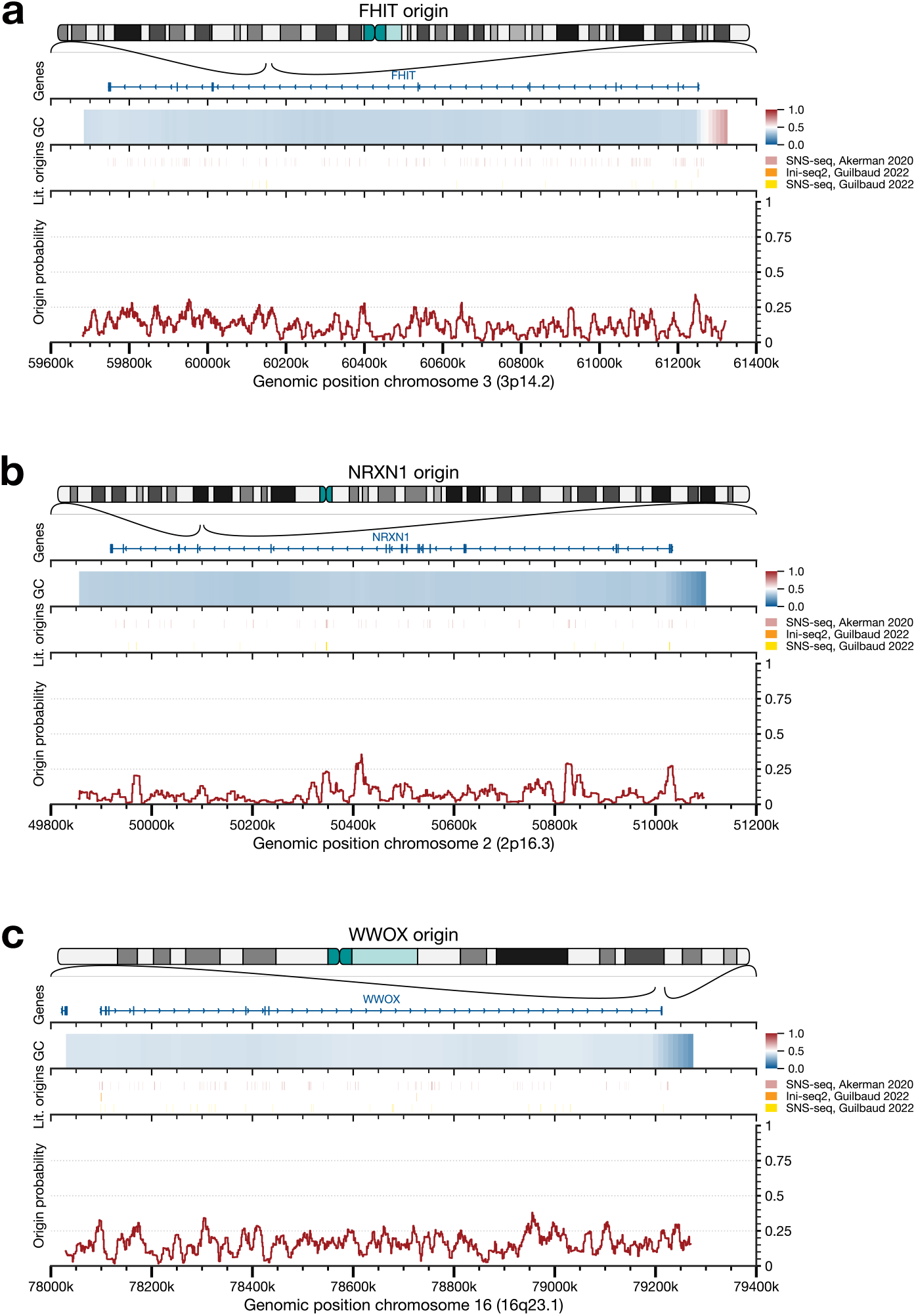
ORILINX predictions in long late-replicating genes. The ORILINX model trained on all human chromosomes was used to predict origin probabilities in 3 regions where no or very few origins were expected based on the literature [24]. **a)** The partially late replicating 1.5 Mb long gene *FHIT*, **b)** the 1.1 Mb long gene *NRXN1* and **c)** the 1.1 Mb long gene *WWOX*.

**Fig. S8.**
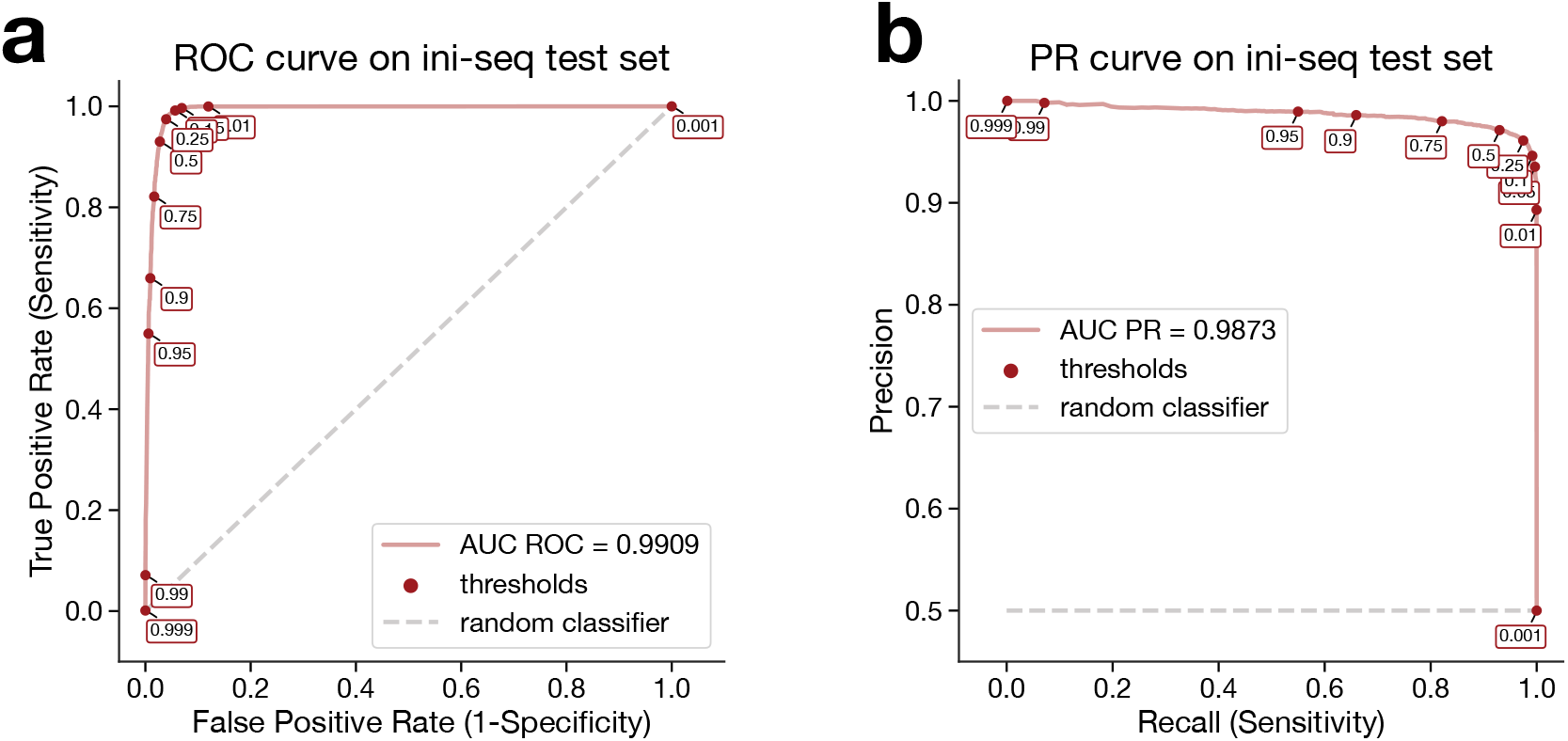
ORILINX version trained on Ini-seq2 data. Following the same training procedure as for the main ORILINX model, we re-trained an ORILINX version using Ini-seq 2 data from the EJ30 cancer cell line [6] for the fine-tuning. The resulting AUC ROC **(a)** and AUC PR **(b)** are presented.

**Fig. S9.**
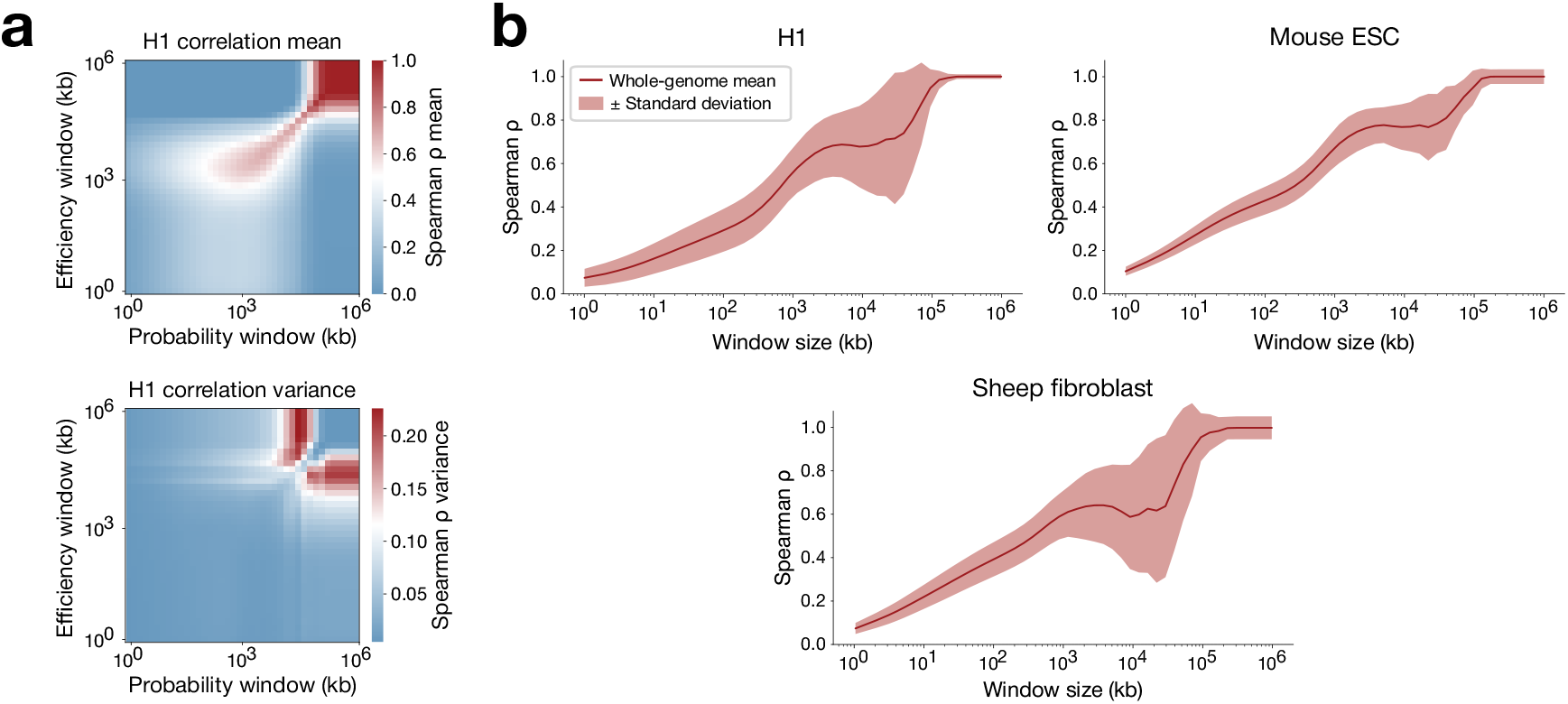
Sensitivity of initiation-efficiency correlations to moving-average resolution. **a)** Heatmaps of Spearman correlation across combinations of probability- and efficiency-smoothing windows in H1, illustrating genome-wide sensitivity to resolution matching. **b)** Dependence of the Spearman correlation on smoothing-window size for multiple species/cell types (H1, mESC, and sheep fibroblast), plotted as mean ± standard deviation across autosomes, highlighting cross-species consistency.

**Fig. S10.**
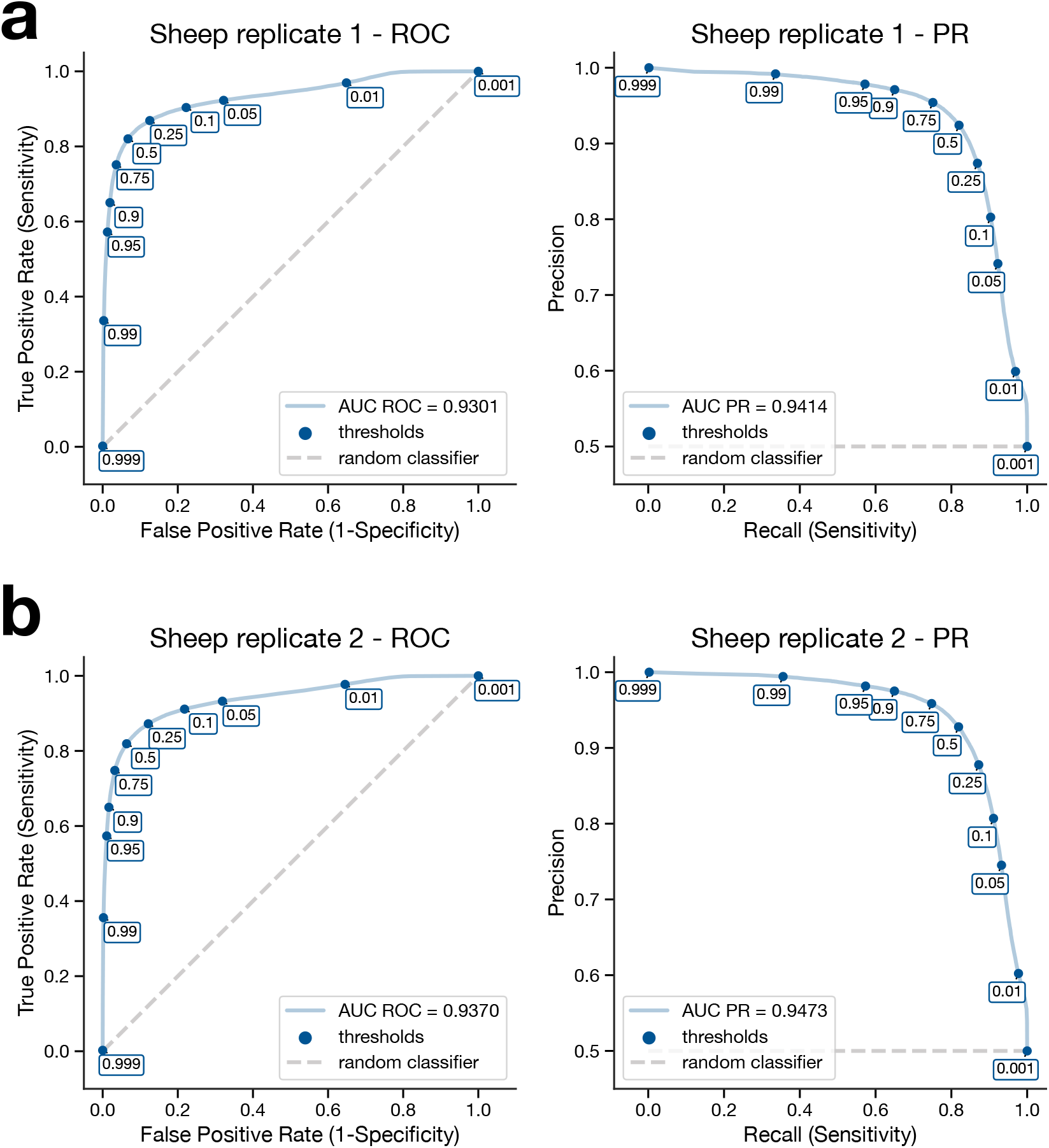
ORILINX predictions across species in individual SNS-seq replicates. AUC ROC and AUC PR curves for individual SNS-seq replicates of the sheep fibroblast cells (see Methods). **a)** Replicate 1 showed an AUC ROC of 0.93 and AUC PR of 0.94. **b)** Replicate 2 showed an AUC ROC of 0.94 and AUC PR of 0.95.

